# Motor imagery practice during arm-immobilization benefits sensorimotor cortical functions and plasticity-related sleep features

**DOI:** 10.1101/828889

**Authors:** Ursula Debarnot, Aurore. A. Perrault, Virginie Sterpenich, Guillaume Legendre, Chieko Huber, Aymeric Guillot, Sophie Schwartz

**Affiliations:** Department of Neuroscience, Faculty of Medicine, University of Geneva, 1211 Geneva, Switzerland; Swiss Center for Affective Science, Campus Biotech, Geneva, 1211 Geneva, Switzerland; Inter-University Laboratory of Human Movement Biology-EA 7424, University Claude Bernard Lyon 1, Villeurbanne, France; Sleep, Cognition and Neuroimaging Laboratory, Department of Health, Kinesiology and Applied Physiology, Concordia University, Montreal, Canada

**Keywords:** motor imagery, sensorimotor representation, cortical excitability, sleep, arm-immobilization

## Abstract

Motor imagery (MI) is known to engage motor networks and could compensate for the maladaptive neuroplasticity elicited by immobilization. This hypothesis and associated underlying neural mechanisms remain underexplored. Here, we investigated how MI practice during 11 h of arm-immobilization influences sensorimotor and cortical representations of the hands, as well as sleep. Fourteen participants were first tested after a normal day, followed by two 11-h periods of immobilization, either with concomitant MI treatment or control tasks. Data revealed that MI prevented the consequences of immobilization: (i) alteration of the sensorimotor representation of hands, (ii) decrease of cortical excitability over the primary motor cortex (M1) contralateral to arm-immobilization, and (iii) reduction of sleep spindles over both M1s. Furthermore, (iv) the time spent in REM sleep was significantly longer after MI. These results support that implementing MI during immobilization can limit the deleterious effects of limb disuse, at several levels of sensorimotor functioning.

---

A period of immobilization does not only impair musculoskeletal motor functions, but elicits substantial modifications at the brain level (Furlan et al., 2016). Specifically, short-term immobilization of the upper-limb (10 h to 24 h) may modulate the excitability of the primary motor cortex (M1) (Avanzino et al., 2011), reorganize sensorimotor functions (Burianova et al., 2016), alter the sensorimotor representation of the non-used hand (Meugnot et al., 2014, Debarnot et al., 2018), and affect sleep characteristics (i.e. spindles and slow waves, Huber et al., 2006). Previous research established that ensuring a stream of sensory inputs during immobilization may protect against motor degradation (Lissek et al., 2009, Avanzino et al., 2014). In particular, recent data showed that proprioceptive vibrations (80 Hz) delivered onto the immobilized arm contributed to prevent the depression of excitability over contralateral M1 (Avanzino et al., 2014). Based on this finding, here we postulated that comparable benefits could be induced by motor imagery practice (MI). MI typically involves the mental rehearsal of an action through visual and proprioceptive inputs without any overt body movement (Jeannerod and Decety, 1995). In the clinical context, accumulating evidence suggests that MI is a reliable, cost-effective and easily implementable substitute for executed movements: MI activates motor networks, which in turn can promote functional recovery (Lotze and Halsband, 2006, Sharma et al., 2006). In case of experimental upper limb immobilization, however, contrasting results on MI efficiency have precluded any clear-cut conclusion. On the one hand, MI practice during arm-immobilization was found to attenuate strength loss (Clark et al., 2014), and to prevent the impairment of both movement preparation time (Stenekes et al., 2009) and sensorimotor representation of the immobilized limb (Meugnot et al., 2015). On the other hand, Crew et al. (2006) and Bassolino et al. (2014) failed to show that MI was effective in compensating the maladaptive plasticity induced by limb disuse. One main aim of the present study was precisely to further clarify whether and how MI practice may alleviate deleterious effects caused by short-term immobilization.

The effects of short-term immobilization have also been studied and observed in the sleeping brain. Notably, following one day with 12 h of immobilization, Huber et al. (2006) observed a reduction of slow wave activity (1-4 Hz) and spindles activity (13-14 Hz) over M1 contralateral to the immobilized arm, predominating during the first NREM sleep period (< 1 h). The authors proposed that these local changes in sleep oscillations after immobilization may be linked to brain plasticity processes, mirroring those induced after motor learning (Huber et al., 2007, Huber et al., 2006, Huber et al., 2004). While it has been established that motor consolidation also occurred after MI, much like for actual motor practice, and yielded subsequent motor improvement (Debarnot et al., 2011a, Debarnot et al., 2009), changes in plasticity-related sleep features following MI are still unknown. Debarnot et al. (2011b) emphasized the importance of NREM sleep for effective MI consolidation, whereas Speth and Speth (2016) underlined the key role of REM sleep. Therefore, experimental limb immobilization does not only provide a useful model for clinical conditions associated with limb disuse (e.g. orthopedic injury, stroke), it may also help to uncover the effects of daytime MI practice on motor-related plasticity mechanisms during wakefulness and during subsequent sleep.

In the present study, we used a randomized crossover within-subjects design to investigate whether MI training during arm-immobilization could counteract the adverse behavioral and neural effects observed after the immobilization period, as well as during subsequent sleep. We expected that MI practice during arm-immobilization would (i) prevent the decrement in the sensorimotor representation of the immobilized limb, (ii) preserve cortical excitability over M1 regions, and (iii) normalize sleep features.

## RESULTS

Fourteen healthy participants (mean age ± SD: 25.43 ± 2.56, nine women) were tested after three different daytime conditions, each separ ted by 1 week (Figure 1). All participants first spent 11 h of normal daily routine (i.e. NOIMMO condition). Then, they underwent the two remaining conditions with their right, dominant arm immobilized during 11 h, either with concomitant MI treatment (15 min each two hours; IMMO+MI condition) or without MI but a cognitive control task (i.e. dice game), with the same time schedule (IMMO–MI condition). The order of the IMMO+MI and IMMO–MI conditions was pseudo-randomized across participants. At the end of each of the 11-h period (NOIMMO, IMMO+MI, IMMO–MI), we tested all participants for their sensorimotor representations of both hands using a mental rotation task (i.e. hand laterality judgment task), cortical excitability for both M1s by means of transcranial magnetic stimulation (TMS), and sleep features using polysomnography (PSG) recordings.

**Figure 1:**
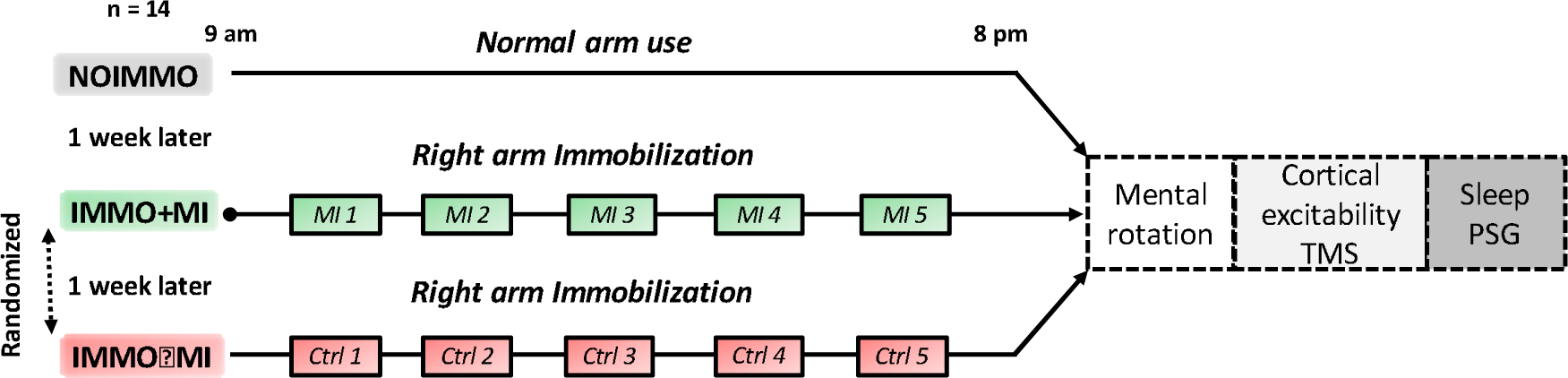
Experimental design. Participants underwent three experimental conditions, separated by one week each, including one normal 11-h awake period without immobilization (NOIMMO), 11 h of arm immobilization with MI treatment (IMMO+MI condition; including 5 MI sessions), or a similar immobilization period without MI (IMMO–MI condition; including 5 control task sessions). After each experimental condition, we assessed changes in sensorimotor representations (mental rotation task) and cortical excitability (TMS) over both M1s. PSG data were recorded during the night following each daytime condition.

### Preliminary note

The mean score of the Stanford Sleepiness Scale assessed at 8 pm (i.e. before starting the multimodal assessment) did not differ as a function of conditions (mean ± SD; NOIMMO: 1.92 ± .64, IMMO+MI: 1.83 ± .55, IMMO–MI: 2.23 ± .89). Actigraphy data confirmed that all participants complied with the instructions regarding the right-arm immobilization procedure. A repeated measures ANOVA on the MET (i.e. metabolic equivalent of task) scores revealed main effects of condition (NOIMMO, IMMO+MI, IMMO–MI); F(2, 52) = 37.99, p < .001, η_p_^2^ =.59) and hand (Right, Left; F(1, 26) = 135.29, p < .001, η_p_^2^ = .83), as well as a significant condition x hand interaction (F(2, 52) = 70.21, p < .001, η_p_^2^ = .73). Bonferroni post-hoc tests showed that during the NOIMMO condition, physical activity with the right arm was higher relative to the left (p < .01), in line with the subjects’ right-hand dominance. Importantly, activity of the right immobilized arm significantly decreased in both IMMO+MI and IMMO–MI conditions compared to NOIMMO (all p < .001), and was significantly lower compared to the left arm activity (p < .001 for both IMMO+MI and IMMO–MI conditions). During both IMMO+MI and IMMO–MI conditions, activity of the left arm did not differ from that in NOIMMO (p ≥ .32 for both comparisons), suggesting that the non-immobilized arm was not more used than usual during right-arm disuse. Regarding MI performance across the 5 MI sessions, participants reported that it was rather easy for them to perform the mental exercises (4.15 ± .12 over a maximum score of 5) and to reach vivid imagery (3.87 ± .11 over a maximum of 5; p > .1 for all comparisons between MI sessions).

### Mental rotation tasks

#### Hand laterality judgment

A repeated measures ANOVA performed on the percentage of correct responses did not yield a main effect of condition (F(2,104) = .85, p = .43, η_p_^2^ = .01), but showed a main effect of configuration (two or three dimensional hand stimuli; F(1,52) = 10.21, p < .01, η_p_^2^ = .16), a main effect of laterality (right or left; F(1,52) = 11.87, p < .01, η_p_^2^ = .18), and a significant configuration × laterality interaction (F(1,52) = 4.32, p < .05, η_p_^2^ = .07; see Methods). Post-hoc comparisons revealed that performance was better for the 3D (95.31 ± 1.81) compared to the 2D (92.11 ± 1.99) configurations, and that left hand stimuli (96.14 ± 1.71) were more correctly identified compared to the right hand ones (91.96 ± 2.09).

A similar ANOVA on response times yielded main effects configuration (F(1,52) = 4.54, p < .05, η_p_^2^ = .08) and condition (F(2,104) = 24.05, p < .001, η_p_^2^ = .32), but no effect of laterality (F(1,52) = .45, p = .50, η_p_^2^ = .01), and no interaction. Bonferroni post-hoc tests revealed that participants were faster at recognizing 3D (vs. 2D; p < .05) configurations, and also both left and right hand stimuli after the IMMO+MI condition compared to NOIMMO and IMMO–MI conditions (p < .001 for each condition comparison with IMMO+MI; Figure 2), while there was no difference between NOIMMO and IMMO–MI (p = .69). These results support that immobilization impacted hand representations after the IMMO–MI condition (i.e. no improvement relative to the NOIMMO, the latter being always administered during the first visit), while MI treatment during arm-immobilization improved hand representations.

**Figure 2:**
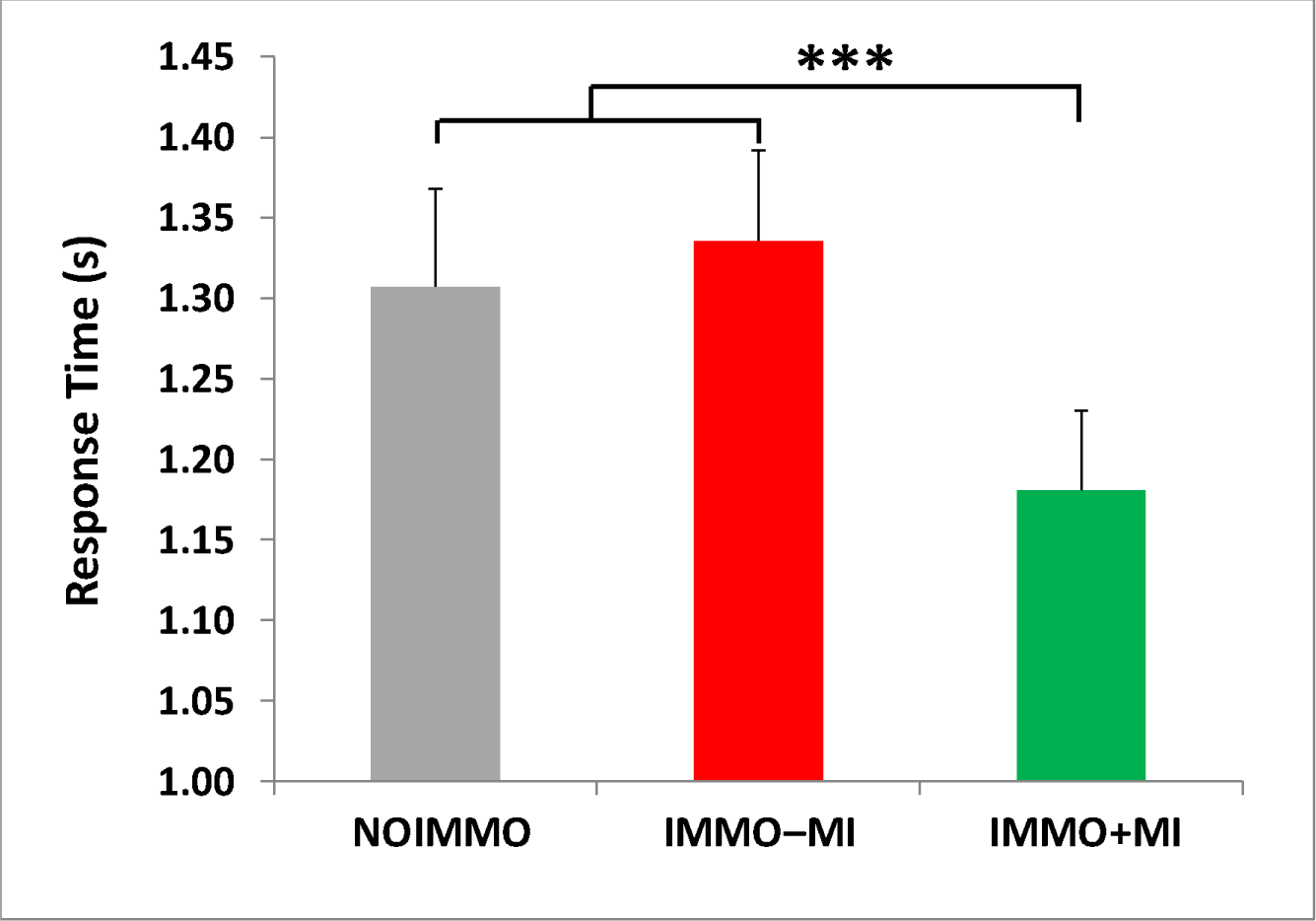
Response times during the hand laterality judgment task. Independently of hand laterality, response times after the IMMO+MI condition was significantly faster compared to NOIMMO and IMMO–MI conditions. Error bars indicate the standard error of the mean. *** p < .001.

#### Alphanumeric normal-mirror judgment task (control task)

A repeated measures ANOVA on the percentage of correct responses for the control mental rotation task revealed a significant effect of configuration (F(1,26) = 48.71, p < .0001, η_p_^2^ = .65) because participants were more accurate for the normal (94%) rather than the mirror (72%) configuration of stimuli. The ANOVA also yielded a condition x configuration interaction (F(2,52) = 3.88.71, p < .0001, η_p_^2^ = .65), and post-hoc analyses showed that normal stimuli were better identified after the IMMO+MI compared to the NOIMMO (p < .05), but not compared to the IMMO–MI condition (p = .20), while performance on mirror stimuli did not show such a modulation by IMMO+MI condition (p < .08 vs. NOIMMO and IMMO–MI conditions).

A similar ANOVA on response times showed a main effect of configuration (F(1,26) = 4.55, p < .05, η_p_^2^ = .15), as well as a main effect of condition (F(2,52) = 4.36, p < .01, η_p_^2^ = .14), but no condition x configuration interaction (F(2,52) = .28, p = .75, η_p_^2^ = .01). Post-hoc tests indicated faster responses for normal (vs. mirror) configurations in both IMMO+MI and IMMO–MI conditions compared to the NOIMMO (p < .01 and p < .05, respectively).

### Transcranial magnetic stimulation

One participant was excluded from TMS analysis due to very high resting motor threshold (RMT > 80% machine output).

Repeated measures ANOVA on the resting motor threshold (RMT) in both M1s showed a main condition effect (F(2,48) = 35.90, p < .01, η_p_^2^ = .15), and post-hoc analyses revealed a lower RMT in the NOIMMO condition over the left M1 compared to the IMMO+MI (p < .05; Supp. Table 1). Comparing the motor evoked potential (MEP), obtained by means of recruitment curve, between conditions (Figure 3A and 3B) revealed a main effect of intensity (F(4,120) = 49.73, p < .001, ηp^2^ = .62), as well as a significant condition x laterality interaction (F(2,240) = 21.49, p < .001, ηp^2^ = .15). Post-hoc analyses showed that MEP amplitude recorded after the IMMO–MI condition was significantly lower than after NOIMMO (p < .001) and IMMO+MI (p < .01) conditions. Most importantly, ANOVA performed on MEP slope for both M1s showed a condition x laterality interaction (F(2,48) = 3.05, p = .05, η_p_^2^ = .11). Post-hoc tests indicated that immobilization reduced the slope of left M1 after the IMMO– MI condition relative to NOIMMO (p <.05), without difference when comparing the IMMO+MI and IMMO–MI conditions (Figure 3C and 3D).

**Figure 3:**
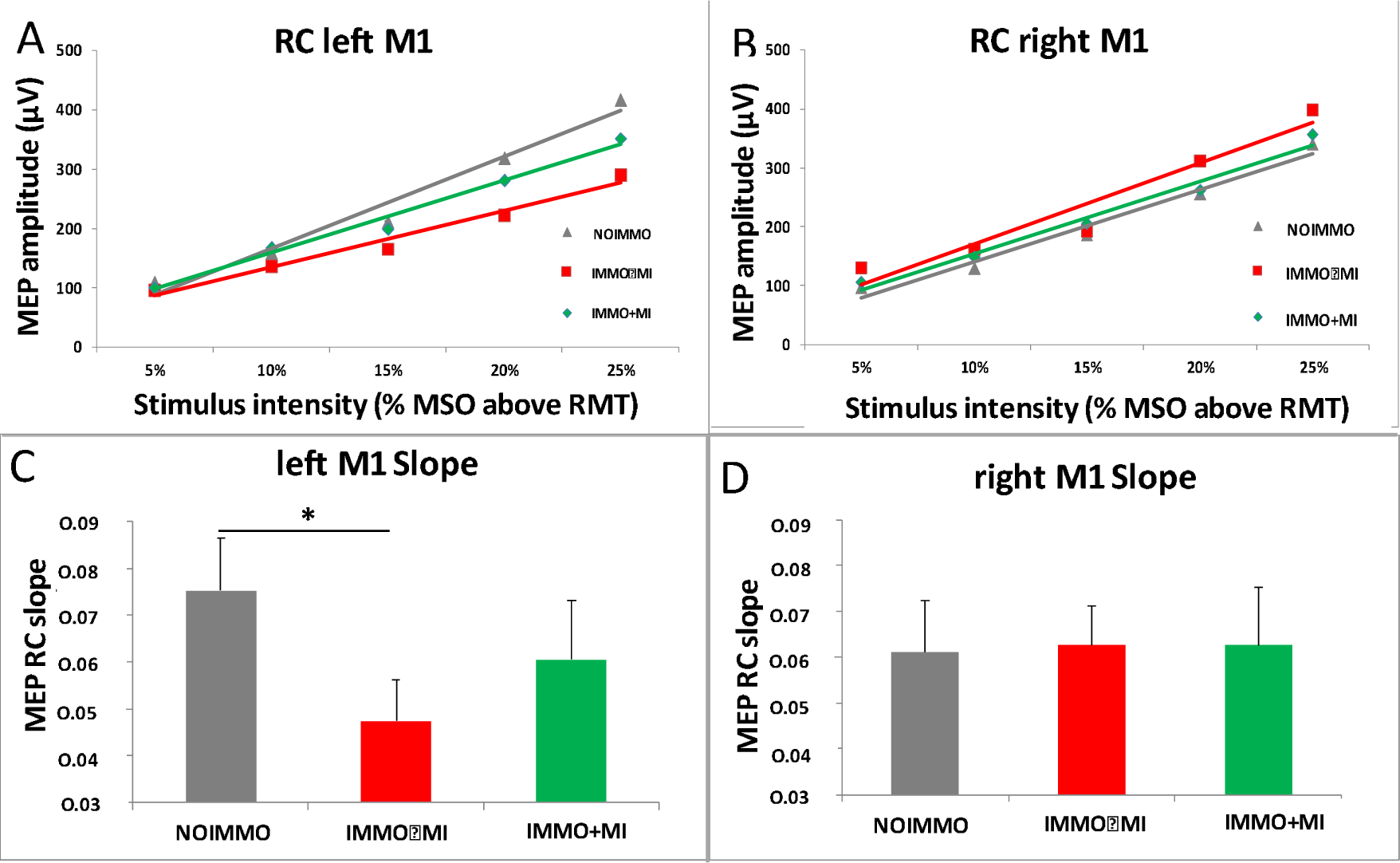
Cortical excitability over left and right M1 after NOIMMO, IMMO–MI and IMMO+MI conditions. (A) MEP amplitude from the left M1 at increasing strength of MSO intensity for the three conditions. (B) Slope data showed a significant reduction in the cortical excitability over left M1 after immobilization alone (IMMO–MI) relative to the NOIMMO condition, while MI practice limited the deleterious effect of immobilization. (C) MEP amplitudes over right M1. (D) Slope data showed no significant differences over the right M1 as a function of conditions. MEP, motor evoked potential; MSO, maximum stimulator output; M1, primary motor cortex; RC, recruitment curve; RMT, resting motor threshold.

We used spearman’s correlation analyses to assess the possible relationship between changes in the corticospinal excitability (i.e. MEP slope) with those at the sensorimotor representation level (i.e. RTs in the mental rotation task), but found no significant correlation regarding the influence of MI treatment [IMMO+MI – IMMO–MI], or of immobilization [NOIMMO – IMMO–MI].

### Polysomographic data

#### Sleep Architecture

Participants showed overall normal sleep patterns across the three PSG nights (Figure 4A and Supp. Table 2). We first tested whether experimental conditions affected the sleep architecture of the whole night recordings. An ANOVA on stage duration (min) with sleep stage (N1, N2, N3, REM) and Condition (NOIMMO, IMMO+MI, IMMO–MI) as within-subjects factors showed an main effect of sleep stage (F(3, 117) = 329.92 ; p < .001, *η*_*p*_ ^2^ = .89), no overall effect of Condition (F(2, 39) = .73 ; p = .48, *η*_*p*_ ^2^ = .03), or interaction (F(6, 117) = .75; p = 60). A similar pattern of results was obtained using percentages of the total sleep period (TSP), while no other change in sleep latencies, total sleep time [TST], and sleep efficiency, was found when comparing experimental conditions (all p > .05).

**Figure 4:**
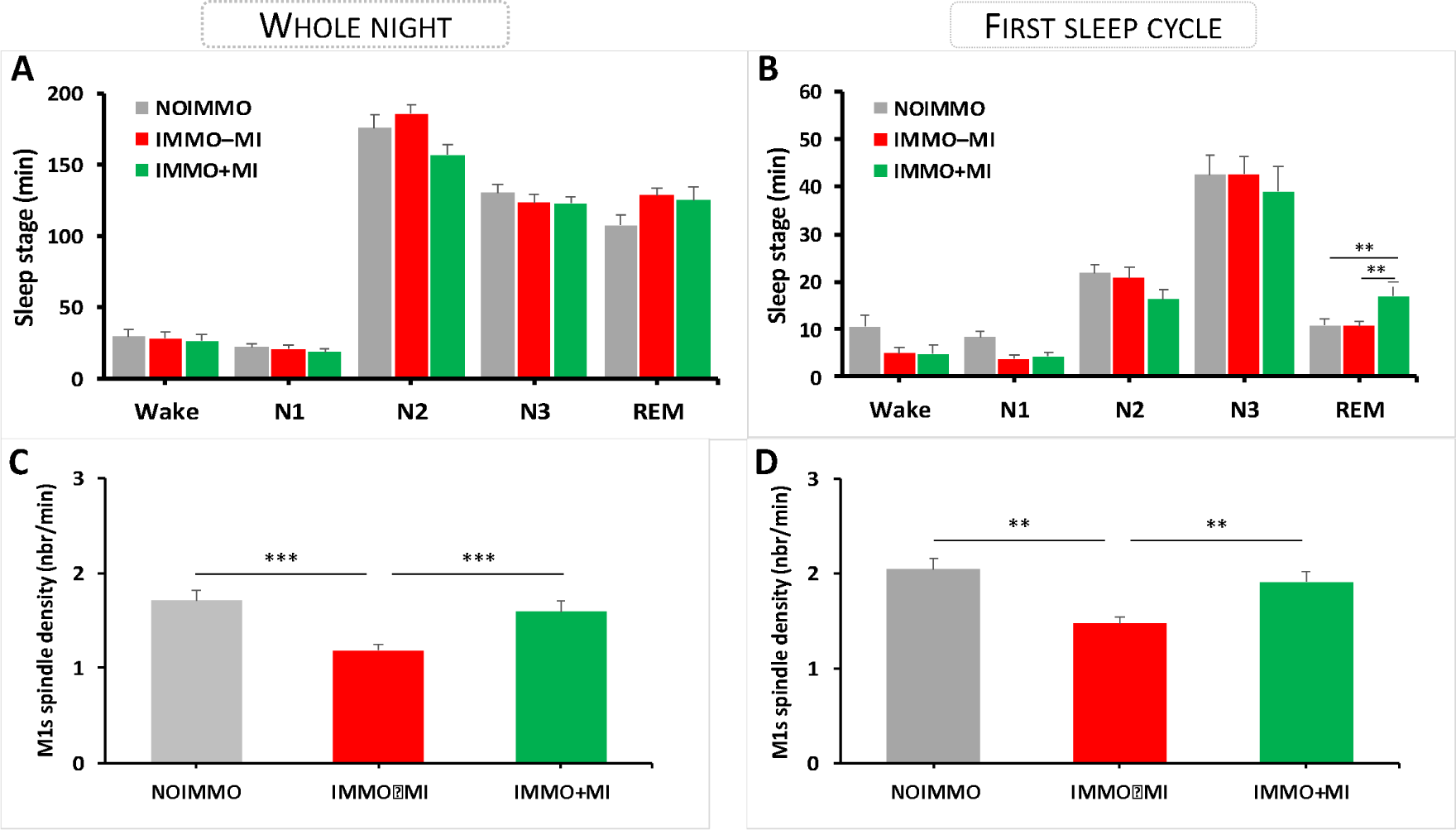
Sleep architecture and spindles density over both M1s during the whole night and the first sleep cycle after each condition (NOIMMO, IMMO+MI, IMMO–MI). (A) Sleep stage duration (min) for each condition during the whole night, and (B) during the first sleep stage. REM sleep duration increased in the IMMO+MI condition relative to NOIMMO and IMMO– MI. (C) Reduced spindle density (nbr/min) over the left and right M1 in the IMMO–MI compared to NOIMMO and IMMO+MI conditions during the whole night, and (D) during the first sleep cycle.

Because studies investigating the impact of learning on subsequent sleep reported effects predominating early in the night sleep, most often during the first sleep cycle (Huber et al., 2006, Huber et al., 2004), we performed the same analyses on the first sleep cycle (Figure 4B). Regarding stage duration (min), we again found the expected main effect of sleep stage (F(3, 117) = 130.38; p < .001, *η*_*p*_ ^2^ = .77), no effect of Condition (F(2, 39) = .99; p = .37, *η*_*p*_ ^2^ = .04), but a significant interaction between sleep stage and Condition (F(6, 117) = 2.35; p = .03). This effect was due to an increase in the time spent in REM after MI treatment (17 ± 2.97 min), as compared to NOIMMO (10.75 ± 1.80 min, p = .01) and to IMMO–MI (10.57 ± .96 min, p = .01). No modulation by MI conditions arose for other sleep parameters (see above, all p > .05).

We further examined the possible links between changes in the time spent in REM (min) during the first sleep cycle with the corticospinal excitability (i.e. MEP slope), and the sensorimotor representation of hands (i.e. RTs), and found no significant correlation regarding the effect of MI treatment [IMMO+MI – IMMO–MI], or of immobilization [NOIMMO – IMMO–MI].

#### Sleep spindles during NREM sleep

Because sleep spindles are known to be modulated by prior learning and experience (Boutin et al., 2018, Morin et al., 2008), including local effects after motor adaptation and limb immobilization (Hoedlmoser et al., 2015, Huber et al., 2006), we tested whether spindles may also be affected in our experiment. We first looked at the total number of spindles detected over left and right M1 during NREM sleep (N2 + N3) from the whole sleep night (Supp. Table 3). An ANOVA with electrode location (left M1, right M1) and condition (NOIMMO, IMMO+MI, IMMO–MI) as within-subjects factors revealed a main effect of condition (F(2, 40) = 18.73; p < .001, *η*_*p*_ ^2^ = .48), but no effect of electrode location (F(1, 20) = .29; p = .59, *η*_*p*_ ^2^ = .01), or interaction (F(2, 40) = .51; p = .60, *η*_*p*_ ^2^ = .02). To clarify the main effect of condition, we performed Bonferroni post-hoc comparisons and showed that compared to the number of spindles was lower during IMMO–MI condition (372.79 ± 24.32; p < .001) compared to NOIMMO (519.66 ± 17.61) and IMMO+MI conditions (431.17; p < .01). Likewise, an ANOVA on spindle density measures revealed a main effect of condition (F(2, 40) = 15.86; p < .001, *η*_*p*_ ^2^ = .44), no effect of electrode location (F(1, 20) = .23; p = .63, *η*_*p*_ ^2^ = .01) and no interaction, F(2, 40) = 1.02; p = .36, *η*_*p*_ ^2^ = .04). Bonferroni post-hoc tests showed a decreased density of spindles after the IMMO–MI condition relative to the NOIMMO and IMMO+MI condition (all comparison p < .001; Figure 4C). Noteworthy, there was no difference in the density of spindles between NOIMMO and IMMO+MI condition over both M1s (p = .70).

Huber et al. (2006) reported that spindle activity was locally modulated (here decreased) following 12 h of arm-immobilization (Huber et al., 2006) and, in line with several studies looking at the effects of prior learning on sleep spindles (Morin et al., 2008, Bang et al., 2014), they found that this modulation occurred at the beginning of the sleep period. Thus, and like we did for the sleep architecture above, we analyzed the number and density of sleep spindles during the first NREM period, again distinguishing between electrodes over the left and right M1 (Supp. Table 4). For the total number of sleep spindles (during N2 + N3), we found a significant main effect of condition (F(2, 40) = 17.07; p < .001, *η*_*p*_ ^2^ = .46), no effect of electrode location (F(1, 20) = .31; p = .58, *η*_*p*_ ^2^ = .01) or interaction (F(2, 40) = .66; p = .52, *η*_*p*_ ^2^ = .03). We observed similar effects for density measures (condition, F(2, 40) = 12.30; p < .001, *η*_*p*_ ^2^ = .38; electrode location, F(1, 20) = .21; p = .64, *η*_*p*_ ^2^ = .01; interaction, F(2, 40) = 1.20; p = .31, *η*_*p*_ ^2^ = .05). Bonferroni post-hoc tests revealed a decreased number of spindles after the IMMO–MI condition relative to the NOIMMO and IMMO+MI condition for both the number and density of spindles (all comparison p < .01; Figure 4D). Importantly, there was no difference in the number or density of sleep spindles over both M1s between NOIMMO and IMMO+MI condition (p = .17 and p = .85, respectively). We further analysed the potential relationship between changes in the density of sleep spindles (whole night and first sleep cycle) with the corticospinal excitability (i.e. MEP slope), and the sensorimotor representation of hands (i.e. RTs). No significant correlations arose from these analyses.

#### EEG Spectral analysis

Two separate ANOVAs on the whole night and first sleep cycle were performed with frequency bands data (slow oscillations, delta, theta, alpha, sigma, spindle power range, and beta), electrode location (left M1, right M1) and condition (NOIMMO, IMMO+MI, IMMO–MI) as within-subjects factors. All analyses yielded a main effect of Frequency bands (all p < .001, as expected for different sleep stages), but no effect of condition (all p ≥ .26), electrode location (all p ≥ .57), or interaction (all p ≥ .82).

## DISCUSSION

We designed the present study to investigate the effects of MI practice administered during 11 h of arm-immobilization on the sensorimotor and cortical representations of both hands, as well as on sleep features. We found that MI practice during unilateral immobilization substantially prevented the degradation of sensorimotor representation and spared cortical excitability over left M1 contralateral to the arm-immobilization. Furthermore, the time spent in REM sleep was significantly longer particularly during the 1 ^st^ cycle of sleep following the IMMO+MI condition compared to that after NOIMMO and IMMO–MI conditions. Finally, data revealed that, while immobilization decreased the number of sleep spindles compared to the condition without immobilization, this decrease was no longer significant after the IMMO+MI condition over both M1s.

### Effect of MI practice on the sensorimotor representation of hands

The first important finding of the present study is that mental rotation performance improved after the IMMO+MI but not after the IMMO–MI condition, for both right-and-left hand stimuli. The lack of task-repetition benefit after the IMMO–MI condition for both right and left hand stimuli confirms and extends previous findings by Toussaint et al. (2013) obtained after 48h of non-dominant hand immobilization. Performance gains observed following the IMMO+MI condition further support previous data showing that 15 min of kinesthetic MI (i.e. hand and finger movements using bodily information) performed right before splint removal following 24 hr of left hand immobilization, contributed to faster discrimination of left hand stimuli (Meugnot et al., 2015). In our study, participants were involved in more intensive MI practice (75 min vs. 15 min), with the last MI session performed 1 h before the splint removal, thus supporting that the present MI regime may have not only transiently activated sensorimotor processing, but also promoted its maintenance at a normal functioning level. Interestingly, a recent neuroimaging study showed that 24 h of hand immobilization, without MI practice sessions, decreased the neural activation underpinning MI, specifically over the sensorimotor areas contralateral to the limb disused (Burianova et al., 2016). Taken together, these observations suggest that reactivation of the sensorimotor system by means of MI practice during immobilization contributed to prevent the maladaptive functional consequences of immobilization. This issue is of particular importance in the clinical context where MI is increasingly used to reactivate injured sensorimotor networks to assist the recovery of lost motor functions (Page et al., 2007, Malouin et al., 2013, Di Rienzo et al., 2014).

### Effect of MI practice on corticomotor excitability

A second main result of the present study is that MI prevented the decrease in cortical excitability of left M1 caused by the contralateral arm-immobilization. We found a significant reduction in the slope of RC data after the IMMO–MI condition, plausibly reflecting local synaptic depression induced by unilateral short-term immobilization (Avanzino et al., 2011, Avanzino et al., 2014, Huber et al., 2006, Rosenkranz et al., 2014). Such downregulation of left M1 excitability was not observed after the IMMO+MI condition. This finding challenges data by Bassolino et al. (2014) and Crew et al. (2006) who reported that MI training could not prevent corticomotor depression after upper-limb immobilization. Such discrepancy may be due to differences in the nature and dosage of the MI intervention. Unlike the simple arm/hand movements used in previous studies, here the experimenter guided participants to mentally rehearse movements using different upper-limb joints, i.e., wrist, elbow and shoulder, and the MI program was structured with a regular increment in task complexity, from simple unilateral movements up to bilateral upper-limb movements. Thus, this thorough and progressive MI training design emphasized the voluntary and effortful engagement of complex motor representations. Moreover, instead of an interactive and tailored guidance, previous studies used audiotaped guidelines or visual stimuli displayed on a computer screen. Crucially, the overall time spent to practice MI was significantly longer in the present study (i.e., 75 min vs. 40 min Bassolino et al. and 3 x 30 min across 3 successive days Crews and Kamen). We thus speculate that a relatively intensive MI intervention might be needed to promote significant M1s activation, hence protecting against loss of sensorimotor functions during limb immobilization. This latter assumption is also supported by reports in the motor learning domain showing that MI increases M1 excitability and that such increase is specific to the representation of the limb involved during imagery (Fadiga et al., 1999, Facchini et al., 2002) .

### Effect of MI practice on sleep

A third original aim of the present study was to investigate whether and how MI practice during immobilization influenced the immediately following sleep period. We first observed an increase in the time spent in REM selectively during the first sleep cycle, consistent with sleep changes occurring predominantly early in the night sleep (Huber et al., 2006, Huber et al., 2007, Huber et al., 2004). Motor circuits are known to be activated during REM sleep (Maquet, 2000, Hobson and Pace-Schott, 2002, Braun et al., 1997); this activation may support MI in dreams and the consolidation of motor skills learning (Speth and Speth, 2018, Speth and Speth, 2016, Speth et al., 2015, Smith et al., 2004, Maquet et al., 2000, Li et al., 2017, Stickgold and Walker, 2013, Schwartz and Maquet, 2002, Eckert et al., 2020), although a causal role of REM sleep for motor learning is still debated (King et al., 2017). Following this line of reasoning, here we suggest that practicing MI exercises during arm-immobilization might have potentiated the demand for REM sleep related motor rehearsal and consolidation, yielding longer REM duration. However, whether increased REM sleep after MI induced better consolidation and recovery of motor skills (i.e. motor test after the night sleep) was not tested in the present study and would thus require further investigation. Finally, our result is also consistent with the clinical observation that patients with REM sleep behavior disorders exhibit a large range of motor behaviors, corresponding to enacting dream actions (Leclair-Visonneau et al., 2010, Arnulf, 2010).

As expected, we found that 11 h of immobilization decreased the density of spindles, not only over the contralateral M1, but over both M1s after the IMMO–MI condition (Huber et al., 2006). Importantly, spindle density over motor regions after the IMMO+MI condition was unchanged as compared to NOIMMO. In line with this hypothesis, Debarnot et al. (2011) reported similar MI performance gains after both a short (10 min NREM) and long (NREM and REM) intervening naps (compared to wake period), suggesting that NREM sleep, and particularly spindles activity may have contributed to the MI consolidation process. Altogether, these findings are in accordance with the critical role of sleep spindles in memory reactivation and consolidation during NREM sleep, including for procedural learning (Morin et al., 2008, Boutin et al., 2018).

To conclude, the present results provide strong evidence that MI practice can alleviate the deleterious impact of arm-immobilization on sensorimotor representations, and motor cortex excitability. Our findings also reveal lasting effects of MI practice on subsequent REM sleep (longer REM duration) and NREM sleep (limitation of the reduction in spindle number). Overall, MI contributed to maintain sensorimotor networks activation during immobilization, hence protecting for the maladaptive neuroplasticity occurring during and after immobilization. Based on these observations, we would like to suggest that an early integration of MI in the context of limb disuse might help to promote recovery of lost motor functions (e.g. during classical physical therapy after orthopaedic intervention). Moreover, MI is increasingly used to control external device such as in electroencephalographic-based brain computer interfaces (Hamedi et al., 2016). Early MI practice after immobilization or in post-stroke rehabilitation programs might help to preserve the cortical excitability over M1 contralateral to the limb disused, and thereby increase the opportunity for the patient to benefit from brain computer interface-based interventions.

## METHODS

### Participants

To be eligible for the study, the fourteen healthy participants had to comply with the following criteria: (1) be right-handed according to the Edinburgh inventory (score >70, Oldfield, 1971); (2) have a Pittsburg Sleep Quality Index < 5 (Buysse et al., 1989) ; (3) have an intermediate circadian typology score from 42 to 58 (Horne and Ostberg, 1976); (4) have an Epworth Sleepiness Scale score < 11; (5) have a Beck Depression inventory score < 9; (6) have a total MI ability score on the Movement Imagery Questionnaire ≥ 63 (corresponding to 75% of the maximum score; MIQ-3, Williams et al., 2012), and (7) have no previous history of orthopedic problems for the right hand and arm. Exclusion criteria included (1) the presence of ferromagnetic metallic implants or a pacemaker; (2) previous neurosurgery, or (3) history of seizures, major head trauma, alcoholism, drug addiction, or any other psychiatric or neurological disorder. All participants were instructed to be alcohol and caffeine free for 24 h prior to and during the experimental days. The protocol was carried out in accordance with ethical standards and was approved by the ethical committee of the Geneva University Hospital. Once the procedure had been fully explained, participants gave written informed consent before inclusion. All participants received an attendance fee.

### Immobilization procedure

During the immobilization conditions, participants were requested to not move the right arm (from the fingers to the shoulder) for 11 h from the morning (9 am) until the beginning of the mental rotation task (8 pm). The duration of the sensorimotor deprivation was chosen based on recent reports demonstrating modifications of M1s excitability (Rosenkranz et al., 2014), with additional local changes during the subsequent night of sleep (Huber et al., 2006). Here, we used a silk glove to reduce the contact between each finger. We further put on a splint, typically used in clinical practice (FERULA, Malleable aluminum thumb-hand right, model “OM6101D/2”), to ensure a complete immobilization of the wrist, as well as the carpometacarpal and metacarpophalangeal joints of all fingers. In addition, a soft shoulder and elbow splint was used to support the forearm in a comfortable way during the 11-h immobilization (DONJOY model IMMO, “DGO GLOBAL”).

To ensure that immobilization was effective, we quantitatively monitored physical activity of both arms during the three experimental conditions, using actimetry (Actigraph GT3X-BT, Pensacola, Florida, USA) on the participants’ left and right forearms. Data were sampled at 30 Hz. Energy cost of physical activity was calculated in metabolic equivalent of task (MET), i.e. one MET equals the resting metabolic rate obtained during quiet sitting (Ainsworth et al., 2000). We also verified that all participants maintained a regular sleep-wake schedule with a minimum of 7 h of sleep per night, at least three days before each experimental day. Compliance to the schedule was assessed using both sleep diaries and wrist actigraphy measures. Finally, every hour during the three experimental days (i.e. 11 times for each experimental day), all participants assessed their current state of alertness by filling out the Stanford Sleepiness Scale (Hoddes et al., 1972).

### Motor imagery sessions

During the IMMO+MI condition, participants performed 5 sessions (one session every two hours, 15 min each) of mental training with the immobilized arm, in a quiet room, without any distracting stimuli in order to help them to focus on MI exercises. The content of MI exercises was varied and included monoarticular and polyarticular movements. Specifically, participants were requested to mentally perform 15 different “monoarticular” movements (e.g. flexion/extension movements of the wrist) with the right-immobilized arm during 1 min for each movement during the two first MI sessions. Then, during the three next MI sessions, they performed five different and complex “polyarticular” movements (e.g. throwing a ball) during 3 min each, i.e. each movement was performed 2 min first and then repeated again during 1 min in order to minimize mental fatigue resulting from maintaining focused attention. For each exercise, the experimenter gave precise instructions about the movement to be performed and asked participants to either sit or stand during MI in order to facilitate the postural congruency related to the movement (e.g. finger tapping was performed in a seated position, performing arm circle was executed while standing up). If needed, participants could first practice the movement with the non-immobilized arm prior to MI rehearsal with the immobilized arm (Toussaint and Blandin, 2010). They were asked to imagine themselves performing movements using a combination of visual and kinaesthetic imagery, i.e. imagining movement within one’s own body and perceiving the induced proprioceptive sensations. They were instructed to say “go” when they started the mental simulation of movement with the immobilized arm. The experimenter watched the participant continuously in order to check that no physical movement with the immobilized arm was produced during MI, and used a stopwatch to indicate the end of an exercise to the participant. After each exercise, participants were asked to rate the difficulty and the quality of their mental images using two Liker-type scales (from 1 = very difficult/poor mental representation to 5 = very easy/vivid mental representation).

### Post-immobilization/NOIMMO assessment of sensorimotor functions

#### Mental rotation tasks

After the 11-h immobilization or NOIMMO period, participants were administered two mental rotation tasks, i.e. a hand laterality judgment task and an alphanumeric normal-mirror judgment task (control task). They sat on a chair at a distance of 50 cm from a 17-inch computer screen with their hands on their knees. They were in a quiet room, without any distracting stimuli, in order to help them to focus on the task. In the hand laterality judgment task, participants were asked to identify whether a hand stimulus presented visually on the screen was depicting a left or right hand. Hand stimuli were depicting either simple, two dimensional views (dorsum or palm view with four possible orientations: 0°, 90°, 180°, or 270°) or more complex, three dimensional views (palm from finger, dorsum from wrist, palm from wrist, or dorsum from finger). In the alphanumeric normal-mirror judgment task, participants had to determine whether the letter “R” or number “2” were presented in a normal or mirror view. Alphanumeric stimuli could be rotated by 0°, 90°, 180°, or 270°. Upright hand and alphanumeric stimuli had 13 cm in height and 7 cm in width.

At the beginning of each trial, a fixation cross was displayed (0.5 s) in the center of the screen. Then, the stimulus was presented and remained visible during 2.5 s. Participants gave a verbal response (left/right in the hand mental rotation task; yes/no in the alphanumeric mental rotation task for normal or mirror image, respectively). Verbal responses were recorded using a microphone connected to the computer. Response time was defined by the time from stimulus onset to the beginning of the audio signal corresponding to the participants’ response. For each trial, the experimenter also collected response accuracy manually.

During the NOIMMO condition, all participants familiarized with the hand laterality judgment task during 6 practice trials of hand stimuli. Right after, the mental rotation test included four randomized blocks of hand laterality judgment task (two blocks of two dimensional, and two blocks of three dimensional items). Each block contained 16 trials with 8 left and 8 right hand pictures, presented in a random order. Then, participants familiarized with the alphanumeric normal-mirror judgment task with 6 practice trials of alphanumeric stimuli (three letters and three numbers) before performing two blocks of 16 trials with 8 normal and 8 mirror alphanumeric items.

The hand mental rotation task was always performed before the alphanumeric mental rotation task, as Toussaint and Meugnot (2013) demonstrated that the effects induced by arm immobilization may be reduced when participants first perform a non-body mental rotation task. During each condition (NOIMMO, IMMO+MI and IMMO–MI), participants thus completed a total of 64 trials of hand laterality judgment task followed by 32 trials of alphanumeric normal-mirror judgment task. During the IMMO+MI and IMMO–MI conditions, participants were no longer familiarized with the tests.

Stimulus presentation, data recording and analysis were handled by a homemade MATLAB program (The MathWorks, Natick, MA, USA), incorporating the Psychtoolbox (Brainard, 1997).

#### Transcranial Magnetic Stimulation

We used TMS to assess cortical excitability of left M1 (contralateral to the immobilized-arm) and then right M1 (ipsilateral to the immobilized-arm) by means of resting motor threshold (RMT) and recruitment curve (RC) after each condition. A figure-of-eight coil with wing diameters of 70 mm, connected to a Magstim 200 magnetic stimulator (Magstim, Whitland, Dyfed, UK) was placed tangentially on the scalp over M1; the handle pointed backward and laterally at a 45° angle to the sagittal plane inducing a posteroanterior current in the brain. First, we identified the optimal spot for eliciting motor-evoked potentials (MEPs) in the right first dorsal interosseous (FDI) muscle and the location was marked on an EEG cap fitted on the participant’s head to provide a reference landmark for each TMS session. Additionally, motor cortices marks over the cap was duplicated on the scalp of participants to attach EEG electrodes for subsequent sleep polysomnographic recording. The RMT was defined as the minimal TMS intensity required to evoke MEPs of about 50 μV peak-to-peak in the targeted muscle in 5 out of 10 consecutive trials and was expressed as a percentage of maximum stimulator output (MSO). Once we found the RMT over left M1, RC was assessed by measuring peak-to-peak amplitude (expressed in μV) of MEPs elicited at stimulus intensities of 5%, 10%, 15%, 20%, and 25% of MSO above individual RMT (obtained on each experimental day). Five trials were recorded at each intensity, and MEP amplitude was then averaged. The same RMT and RC procedure was then performed over right M1.

#### Electromyography recording

Electromyography (EMG) was recorded and monitored continuously on-line during each TMS session. EMG recordings used the following montage: active electrodes were attached to the skin overlying first dorsal interosseous muscle, reference electrodes were placed over the first metacarpophalangeal joints and ground electrodes were placed over the wrist bone. Acknowledge 4 software along with MP36 unit (Biopac systems, Goleta, CA, USA) were used to acquire EMG signal. The EMG signals were amplified and filtered online (20 Hz to 1 kHz), sampled at a rate of 5000 Hz and stored on a personal computer for display and later offline data analysis.

#### Polysomnographic recording

##### Data acquisition

Polysomnography was acquired in all participants during the night immediately following NOIMMO, IMMO+MI and IMMO–MI conditions, with a V-Amp 16 system (Brain Products, Gilching, Germany), from 11 EEG electrodes (international 10–20 system: LM1, CZ, RM1, F3, FZ, F4, P3, PZ, P4, A1, A2), 3 electrooculogram electrodes, and 2 chin electromyogram electrodes (sampling rate: 250 Hz).

##### Sleep data analysis

Out of the 14 participants, three participants were excluded from spindles and spectral power analysis due to poor EEG signal quality (on one of the nights), while sleep stage scoring was still possible for these participants for all three experimental nights (n = 14). For sleep analyses, we used the PRANA software (version 10.1. Strasbourg, France: Phitools) and custom MATLAB scripts (MATLAB R2009b, Natick, Massachusetts: The MathWorks Inc., 2009). Two scorers blind to the experimental conditions determined the different sleep stages (NREM 1, 2, 3, REM sleep and wake) for each recorded night of sleep on 30-s epochs according to the AASM standards (Medicine., 2007). From the sleep scoring, we computed the total time spent (min) in each sleep stage, as well as the percentage of each sleep stage relative to the total sleep period (TSP; from sleep onset to wake up time) and relative to the total sleep time (TST; TSP minus intra-wake epochs). Sleep efficiency (TST/time in bed*100), and number of intra-awakenings were also calculated. Epochs affected by artefacts were excluded from further quantitative EEG analysis. The EEG spectrum power average was calculated with a 0.2 Hz resolution, by applying a Fast Fourier Transform (FFT; 50% overlapping, 5 s windows, Hanning filter) on the left and right M1 channels. Mean power was calculated for the following frequency bands: slow oscillations (0.4-1 Hz), delta (1-4 Hz), theta (4-8 Hz), alpha (8-12 Hz), sigma or spindle power range (12-15 Hz), and beta (15-30 Hz). The detection of sleep spindles was performed automatically with the PRANA software on each channel following the global standards used in sleep research (Ray et al., 2010, Morin et al., 2008, Dang-Vu et al., 2010): spindles were required to have a duration between 0.5 s and 3 s, an inter-spindle interval no less than 1 s, and a typical, fusiform spindle morphology (waxing and waning amplitude). An experienced scorer visually supervised all detected spindles in each night of each participant.

### Data analysis

For the mental rotation task, we analyzed the percentage of correct responses and mean response time. Only data from correct responses were used to analyze response times. Results obtained on the hand laterality judgment for the three conditions were evaluated by means of ANOVA with condition (NOIMMO, IMMO+MI and IMMO–MI), laterality (left, right) and configuration (two or three dimensional) as within-subjects factors. Likewise, an ANOVA was performed for the alphanumeric normal-mirror judgment task, with condition (NOIMMO, IMMO+MI and IMMO–MI), configuration (normal, mirror) and orientation (0°, 90°, 180°, or 270°) as within-subjects factors.

To evaluate the effects of immobilization on cortical excitability, we compared the resting motor threshold (RMT) using an ANOVA with laterality (left M1, right M1) and condition (NOIMMO, IMMO+MI and IMMO–MI) as within-subject factors. Recruitment curve (RC) values were analyzed by means of an ANOVA with laterality (left M1, right M1), condition (NOIMMO, IMMO+MI and IMMO–MI) and intensity (5%, 10%, 15%, 20%, and 25% of MSO above RMT) as within-subjects factors. We then evaluated the differences between 3 regression slopes using an ANOVA with condition (NOIMMO, IMMO+MI and IMMO–MI) and laterality (left M1 and right M1) as within-subjects factors.

PSG parameters (sleep latencies, N1, N2, N3 and REM data) did not distribute normally (Shapiro-Wilks W test), and were log-transformed prior to statistical analysis. To examine whether daytime conditions (NOIMMO, IMMO+MI and IMMO–MI) influenced sleep architecture, we performed ANOVAs with sleep stage duration (min; N1, N2, N3, REM) and condition (NOIMMO, IMMO+MI and IMMO–MI) as within-subjects factors. Similar ANOVAs were performed with total sleep period (%), sleep latencies and efficiency (min and % respectively). We also analysed spindle features (number, density, duration, peak amplitude and mean frequency) by means of ANOVAs with electrode location (left M1, Right M1) and condition as within-subjects factors. Similar analyses were performed on power spectrum measurements with electrode location and condition as within-subjects factors. All sleep analyses (sleep architecture, spindles features and spectral power data) were conducted separately during whole night of sleep and the first sleep cycle.

We conducted correlation analyses by computing data from the multimodal assessment obtained in each condition to address the effect of MI treatment [IMMO+MI vs IMMO–MI] and the impact of immobilization [NOIMMO vs IMMO–MI]. Spearman’s correlations were performed between the changes observed in sleep architecture (i.e. REM duration) or spindles density, on the one hand, and the sensorimotor representation of hands (i.e. RTs mental rotation task) on the corticospinal excitability (i.e. MEP slope), on the other hand.

Both the Stanford Sleepiness Scale and MET scores for the NOIMMO, IMMO+MI, IMMO–MI conditions were compared using ANOVAs.

Whenever interactions were significant, the origin of the interaction was investigated using planned comparisons with Bonferroni-corrected post-hoc tests. The alpha-level of all analyses was set at *p* < .05.

